# Contrasting adaptations to synaptic physiology of prefrontal cortex interneuron subtypes in a mouse model of binge drinking

**DOI:** 10.1101/2020.01.05.895169

**Authors:** Max E. Joffe, Danny G. Winder, P. Jeffrey Conn

**Affiliations:** Department of Pharmacology, Vanderbilt University, Nashville, TN, 37232, USA.; Vanderbilt Center for Neuroscience Drug Discovery, Nashville, TN, 37232, USA; Vanderbilt Center for Addiction Research, Nashville, TN, 37232, USA; Department of Molecular Physiology and Biophysics, Vanderbilt University, Nashville, TN, 37232, USA

**Keywords:** alcohol, prefrontal cortex, sex differences, synaptic physiology, parvalbumin, somatostatin

## Abstract

Alcohol use disorder (AUD) affects all sexes, however women who develop AUD may be particularly susceptible to cravings and other components of the disease. While many brain regions are involved in AUD etiology, proper function of the prefrontal cortex (PFC) is particularly important for top-down craving management and the moderation of drinking behaviors. Essential regulation of PFC output is provided by local inhibitory interneurons, yet the effects of chronic drinking on interneuron physiology remain poorly understood, particularly in female individuals. To address this gap, we generated fluorescent reporter transgenic mice to label the two major classes of interneuron in deep layer prelimbic PFC, based on expression of parvalbumin (PV-IN) or somatostatin (SST-IN). We then interrogated PV-IN and SST-IN membrane and synaptic physiology in a rodent model of binge drinking. Beginning in late adolescence, mice received 3-4 weeks of intermittent access (IA) ethanol. One day after the last drinking session, adaptations to PV-IN and SST-IN intrinsic physiology were observed in male mice but not in female mice. Furthermore, IA ethanol precipitated diametrically opposing changes to PV-IN synaptic physiology based on sex. IA ethanol decreased excitatory synaptic strength onto PV-INs from female mice and potentiated excitatory transmission onto PV-INs male mice. In contrast, decreased synaptic strength onto SST-INs was observed following IA ethanol in all groups of mice. Together, these findings illustrate novel sex differences in drinking-related PFC pathophysiology. Discovering means to restore PV-IN and SST-IN dysfunction following extended drinking provides opportunities for developing new treatments for all AUD patients.

## **1.** Introduction

Alcohol use disorder (AUD) affects all groups of people (Hasin et al., 2007), however the prevalence of alcohol abuse among women is increasing (Grant et al., 2017) and several concerning findings indicate that women are at high risk for detrimental outcomes. Women are disproportionately affected by consequences of acute intoxication (Gross and Billingham, 1998) and are particularly sensitive to peripheral diseases and cognitive disturbances stemming from chronic alcohol consumption (Nixon et al., 1995; Urbano-Marquez et al., 1995). Furthermore, the prevalence of binge drinking is increasing among women (Grucza et al., 2018) and women who develop AUD progress rapidly through disease milestones (Diehl et al., 2007; Randall et al., 1999), suggesting sex differences regulate the top-down control over drinking. These sex differences are likely mediated, in part, by the medial prefrontal cortex (PFC), a region whose dysfunction is linked with deficits in the ability to control alcohol cravings in AUD patients and animal models (Abernathy et al., 2010; George and Koob, 2010). In addition to this overall relationship, binge drinkers and AUD patients display sex differences in PFC structure (Medina et al., 2008; Squeglia et al., 2012), and women with AUD display opposite patterns of PFC activation during working memory tasks relative to men (Caldwell et al., 2005). Together, these findings provide compelling rationale for continued mechanistic research to understand sex differences in PFC pathophysiology in AUD-like disease models.

Preclinical studies designed to model alcohol-induced changes to PFC function have largely utilized the chronic intermittent ethanol (CIE) exposure paradigm, an animal model of dependence. Using CIE and other chronic treatment models, several labs have described dependence-related changes in PFC physiology to be generally characterized by reduced inhibition and enhanced excitatory synaptic activity (Centanni et al., 2017; Hu et al., 2015; Pava and Woodward, 2014; Pleil et al., 2015; Varodayan et al., 2018). In addition, changes in NMDA receptor function (Hu et al., 2015; Kroener et al., 2012), intrinsic properties (Hu et al., 2015), and PFC network activity (Kroener et al., 2012; Woodward and Pava, 2009), have all been shown to occur as a consequence of long-term alcohol exposure and withdrawal. While the literature clearly demonstrates that PFC pathophysiology develops during dependence, deficits in PFC function are also associated with maladaptive changes in voluntary drinking (Haun et al., 2018; Klenowski et al., 2016; Radke et al., 2017b; Salling et al., 2018; Seif et al., 2013; Siciliano et al., 2019). Moreover, mounting evidence suggests that PFC interneurons may be particularly important for regulating volitional alcohol-seeking. For example, genetic disruption of synaptic transmission on forebrain interneurons, but not projection neurons, decreases drinking (Radke et al., 2017a), and early abstinence from voluntary ethanol specifically increases Fos expression in PFC interneurons (George et al., 2012). While these exciting findings suggest that PFC inhibitory microcircuits instruct drinking moderation, much remains to be learned about the specific cell types and synapses underlying these processes.

A major source of PFC output to the limbic system arises from deep layer pyramidal cells. While pyramidal cells comprise approximately 80% of the neurons in PFC, the remaining interneurons are essential in coordinating the output from the structure (Ferguson and Gao, 2018). Deep layer PFC contains two general classes of local inhibitory interneurons that are readily divided by their form, function, and genetics. The synapses of one class appose the cell bodies of neighboring pyramidal cells, where they exert powerful feedforward inhibition to synchronize PFC network activity and output (Atallah et al., 2012; Sohal et al., 2009). These interneurons, known as “basket cells” or “chandelier cells”, have unique intrinsic properties that can be used to functionally demarcate them from regular-spiking pyramidal cells or low-threshold spiking interneurons. Using this approach, acute ethanol (Woodward and Pava, 2009) and CIE (Trantham-Davidson et al., 2014; Trantham-Davidson et al., 2017) have been shown to disrupt PFC fast-spiking interneuron function. These fast-spiking interneurons exclusively express the Ca^2+^-binding protein parvalbumin (PV). Transgenic mouse lines have been engineered to express fluorescent markers and manipulate protein expression under the control of the PV promotor (Taniguchi et al., 2011), enabling the means to selectively manipulate the physiology of PV-expressing interneurons (PV-INs). Recent studies have leveraged these tools to reveal that GABAA receptor subunit ablation from PV-INs increases binge drinking in male mice, but not female mice, supporting the hypothesis that this interneuron subtype regulates sex differences in top-down control (Melon et al., 2018). In addition to PV-INs, the second class of deep layer PFC interneurons expresses the neuropeptide somatostatin (SST-INs). Relatively little is known about how ethanol regulates SST-IN function, partly because these neurons are more difficult to functionally separate from pyramidal cells without genetic labeling tools. In general, SST-INs project to the superficial dendrites of neighboring pyramidal cells to filter synaptic information as it flows towards the cell body. SST-INs are therefore critical for processing long-range transmission and interactions with subcortical areas (Abbas et al., 2018). In sum, PV-INs and SST-INs serve essential, complimentary functions in mediating feedforward and feedback inhibition in the PFC.

Female rodents exhibit higher levels of voluntary alcohol consumption than male counterparts (Becker and Lopez, 2004; Hwa et al., 2011; Jury et al., 2017; McCall et al., 2013). The behavioral findings are striking and consistent across laboratories, but the mechanisms through which PFC dysregulation contributes to this phenomenon remain unclear. Most previous studies examining PFC dysfunction were conducted in male rodents and interneurons were solely classified based on membrane properties. In addition, most studies examining PFC interneurons were performed following non-contingent exposure, leaving us with a relatively limited understanding of how PFC inhibitory microcircuits adapt following long-term volitional drinking. Based on this, we utilized transgenic fluorescent reporter mice to investigate sex differences in PFC PV-IN and SST-IN physiology in a binge drinking model. After intermittent access (IA) ethanol exposure, we observed intrinsic physiology adaptations in both PV-INs and SST-INs of male mice but not female mice. In PV-INs, IA ethanol generated diametrically opposing changes to synaptic physiology across sexes. By contrast, excitatory synaptic strength onto SST-INs was decreased in all groups of mice after IA ethanol. Collectively, these findings highlight striking and contrasting adaptations to cortical interneuron synaptic physiology induced by long-term voluntary drinking.

## 2. Material and Methods

### 2.1 Mice

Mice were bred and housed in a controlled environment on a standard 12-hour light cycle (on at 6:00 am). Transgenic mice expressing tdTomato fluorescent protein in PFC interneurons were generated by crossing female PV-Cre mice (Jackson Laboratories, Stock No: 017320) or SST-IRES-Cre mice (Jackson Laboratories, Stock No: 028864) with male Rosa26-loxP-STOP-loxP-CAG-tdTomato “Ai9” mice (Jackson Laboratories, Stock No: 007909). All breeding strains were congenic on a C57BL/6J genetic background. Only female PV-Cre mice were used for breeding to mitigate PV-Cre driven recombination that can occur in sperm. All breeding mice were homozygous for the respective transgene, generating heterozygous PV-tdTomato and SST-tdTomato mice suitable for experimentation. Mice were defined as female or male by their external genitalia and the current studies are limited by this definition.

### 2.2 Intermittent access (IA) to ethanol

Mice provided with IA ethanol drink more per day than continuous access controls (Hwa et al., 2011), and the C57BL/6J strain consumes more ethanol than other strains (Belknap et al., 1993; Boyce-Rustay et al., 2008; Rodgers and Mc, 1962). For these reasons, we selected the IA schedule in C57BL/6J mice as a robust rodent model of binge drinking. IA ethanol began during late adolescence (6-7 weeks). For alternating 24-hour periods, mice were provided with free access to ethanol in their home cages. Water and food were always provided *ad libitum*. Mice were individually housed 3-7 days prior to IA ethanol initiation and remained so until sacrifice.

Ethanol was provided 3-4 hours prior to the dark cycle and removed one day later. For the first week of access, the concentration of ethanol was slowly ramped up (3, 6, 10%) to 20% ethanol, which was used for the duration of the study. The amount of ethanol and water consumed per day was measured by weight after each drinking session. As expected, female mice drank more ethanol than matched male mice (Figure 1A and 1B). We observed no sex differences in preference for ethanol over water across the IA access procedure (Figure 1C and 1D).

**Figure 1.**
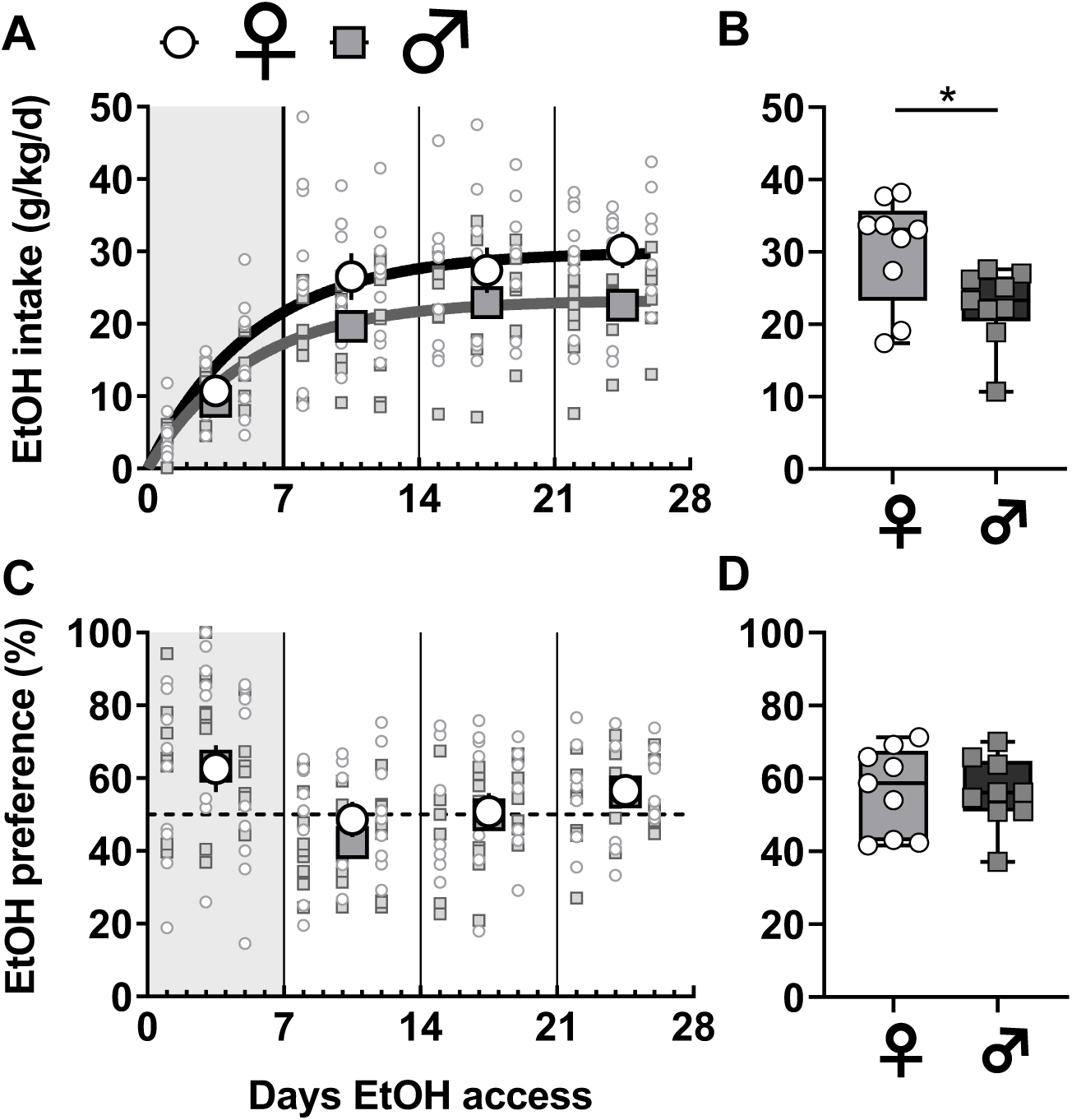
Sex differences in voluntary ethanol consumption under an intermittent access schedule. **(A)** Mice underwent 4 weeks of voluntary drinking during which ethanol was provided every other day on an intermittent access (IA) schedule. The ethanol concentration was ramped during the first week and set at 20% for the duration of the study. Female mice (light circles) displayed increased ethanol intake relative to male mice (dark squares) over the duration of the study (Least squares best-fit, one-phase association: 95% confidence intervals of plateaus {27.1 to 33.6 g/kg; r^2^ = 0.46} vs {21.5 to 25.4 g/kg; r^2^ = 0.53}). N = 9 mice per group. **(B)** Female mice drank more during the last week of the IA ethanol paradigm (30.2 ± 2.5 vs 22.6 ± 1.7 g/kg, *: p < 0.05, t-test). **(C-D)** No sex difference in preference for ethanol over concurrently available water across the IA ethanol paradigm or during the last week of drinking.

### 2.3 Electrophysiology

After 3-4 weeks of 20% ethanol intake, we assessed physiological changes to PV-IN and SST-IN physiology. Mice were sacrificed 16-20 hours after the last session, a time linked to increased PFC interneuron activity (George et al., 2012). Acute prelimbic PFC slices were prepared for whole-cell patch-clamp physiology as described (Di Menna et al., 2018). In brief, mice were anesthetized with isoflurane and decapitated. Brains were rapidly removed without perfusion and submerged in *N*-methyl-D-glucamine solution. Immediately after preparation, coronal slices (300 μM) recovered in warm (30-32 °C) *N*-methyl-D-glucamine solution for 10 minutes and then in room-temperature (22-24 °C) for 1 hour in artificial cerebrospinal fluid, containing (in mM): 119 NaCl, 2.5 KCl, 2.5 CaCl2, 1.3 MgCl2, 1 NaH2PO4, 11 glucose, and 26 NaHCO3. Membrane properties were assessed in current clamp configuration using a potassium-based internal solution (in mM): 125 K-gluconate, 4 NaCl, 10 HEPES, 4 MgATP, 0.3 NaGTP, 10 Tris-phosphocreatine. Cells were dialyzed with internal solution for 5 minutes, after which a series of 20, 1-sec current injections were applied. Injections began at -150 pA, were incremented at 25 pA, and ended at +300 pA. Rm was calculated as the slope of the potential hyperpolarization divided by the injected current. Sag ratio was evaluated based on the resting membrane potential (Vm) and hyperpolarization in response to -150 pA current injection. Sag ratio was calculated as the difference between the peak hyperpolarization and the steady-state, normalized to the steady-state (Joffe et al., 2019). Medium afterhyperpolarization (mAHP) was determined as the magnitude of the decrease in membrane potential during the 0.5-sec period after a current injection resulted in >20 action potentials (generally 200-300 pA for PV-INs and 100-200 pA for SST-INs). Cells were then switched to voltage clamp configuration, and spontaneous excitatory synaptic transmission was collected over 2 minutes. Spontaneous excitatory postsynaptic currents (sEPSCs) were detected with templates specific for each interneuron subtype. Finally, in some cells, the paired-pulse ratio (PPR) of evoked EPSCs was assessed using local electrical stimulation (5-50 µA, 0.1 ms) of superficial layer 5 at 0.2 Hz. For each intersimulus interval (25-400 ms), 6-7 traces were averaged to determine PPR.

### 2.4 Interneuron classification

Neurons were initially selected by tdTomato fluorescence. All PV-tdTomato neurons displayed functional characteristics consistent with fast-spiking interneurons and were included in experiments. While most SST-tdTomato neurons exhibited low-threshold firing consistent with Martinotti cells, approximately one-fourth displayed irregular or fast-spiking-like properties. SST-tdTomato neurons with low Rm (< 150 MΩ), hyperpolarized Vm (< -75 mV), and high rheobase (> 100 pA) were immediately discarded, as they represent ectopic tdTomato expression stemming from transient SST expression during development (Hu et al., 2013), or non-Martinotti type SST-INs (Nigro et al., 2018).

### 2.5 Statistics

The number of cells or mice is denoted by “n” and/or “N” respectively, in each figure legend. Data are generally presented as mean ± standard error or as box plots displaying median, interquartile range, and range. Analyses were performed in GraphPad Prism. Two-tailed Student’s t-test, non-linear least squares best-fit regression, and two-way repeated-measures ANOVA with Bonferonni post-hoc comparisons were used as appropriate. All statistical findings are displayed in the figure legends.

## 3. Results

### 3.1 Basal physiology of PFC PV-INs in female mice and male mice

To assess whether PFC fast-spiking interneurons display sex differences in intrinsic and synaptic physiology, we generated mice expressing tdTomato in PV-expressing neurons (Figure 2A and 2B). Using whole-cell patch-clamp techniques we validated that PV(+) cells represent fast-spiking interneurons in layer 5 prelimbic PFC. Indeed, PV(+) cells display the hallmark characteristics of fast-spiking interneurons (Connors and Gutnick, 1990; Kawaguchi, 1993; Markram et al., 2004; McCormick et al., 1985), including high firing frequency and minimal spike-firing adaptation and hyperpolarization sag (Figure 2B-2F). PV-IN intrinsic properties were comparable between female and male mice, however PV-INs from male mice exhibited a smaller medium afterhyperpolarization (mAHP) than those from female mice (Figure 2G).

**Figure 2.**
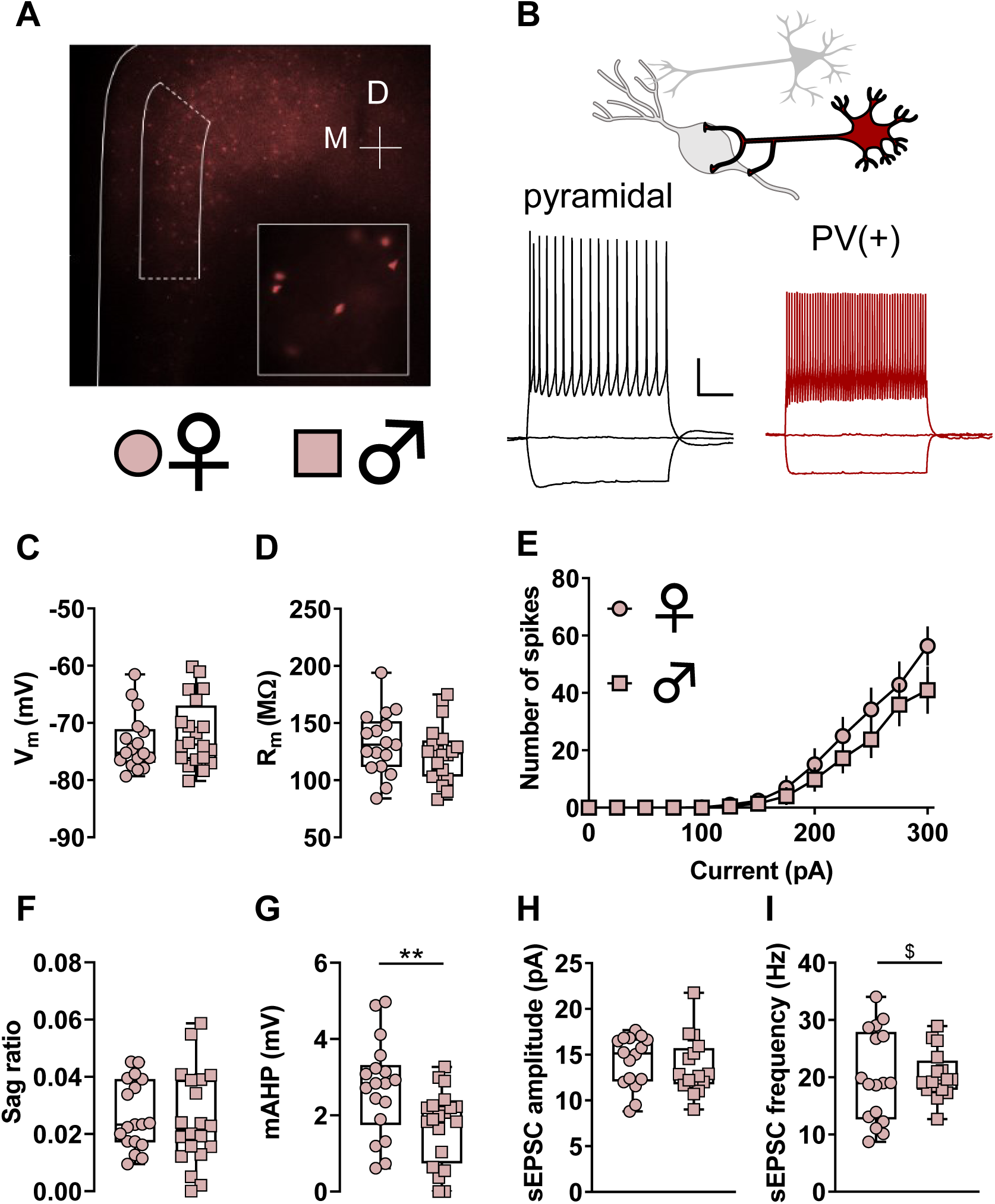
Parvalbumin (PV)-(+) neurons display functional characteristics of fast-spiking interneurons. **(A)** Representative 4X image displaying TdTomato fluorescence in deep layers of the mouse prefrontal cortex. Boxed area indicates prelimbic subregion. Inset, 40X magnification image showing individual PV-expressing interneurons. M, medial; D, dorsal. **(B)** Whole-cell patch-clamp recordings were made from identified neurons in the PFC. Representative current-clamp recordings from an unlabeled pyramidal cell (left, black) and a TdTomato-labeled PV-IN (right, red). The pyramidal cell displays a hyperpolarization-activated sag and accommodating spike firing, physiological features that are minimal or absent in PV-INs. Scale bars indicate 20 mV and 250 ms. **(C)** Resting membrane potential (V_m_) in PV-INs from female mice (circles) and male mice (squares). n/N = 17/9, 20/8 cells / mice. **(D)** No difference in membrane resistance (R_m_) between PV-INs from female and male mice. n/N = 17/9, 20/8. **(E)** No difference in current-evoked spiking between PV-INs from female and male mice. n/N = 16/9, 18/8. **(F)** No difference in hyperpolarization sag ratio between PV-INs from female and male mice. n/N = 18/9, 20/8. **(G)** PV-INs from female mice display greater medium afterhyperpolarization (mAHP) than PV-INs from male mice (2.71 ± 0.30 vs 1.72 ± 0.22 mV, **: p < 0.01, t-test). n/N = 18/9, 20/8. **(H)** No difference in spontaneous excitatory postsynaptic current (sEPSC) amplitude between PV-INs from female and male mice. n/N = 16/9, 16/7. **(I)** No difference in the mean of sEPSC frequency between PV-INs from female and male mice. A difference in the variance of sEPSC frequency was observed between female and male mice (F_16,15_ = 3.526, $: p < 0.01, F test to compare variances). n/N = 17/9, 16/7.

Measurements of basal excitatory synaptic strength were also similar across sexes (Figure 2H and 2I), but the variance of PV-IN sEPSC frequency was greater in female mice (Figure 2I).

### 3.2 Basal physiology of PFC SST-INs in female mice and male mice

To assess the intrinsic and synaptic physiology of PFC low-threshold spiking interneurons, we generated mice expressing tdTomato under control of the SST promotor (Figure 3A and 3B). In layer 5 PFC, most SST(+) cells represent low-threshold spiking interneurons, however some neurons displayed fast-spiking-like phenotypes and were immediately relinquished. Low-threshold spiking SST(+) cells display several characteristics of Martinotti cells (Nigro et al., 2018; Tremblay et al., 2016) including high Rm, depolarized Vm, low rheobase, and firing upon rebound from hyperpolarization (Figure 3B). SST-IN intrinsic properties and measurements of basal synaptic physiology were all comparable across sexes (Figure 3C-3I).

**Figure 3.**
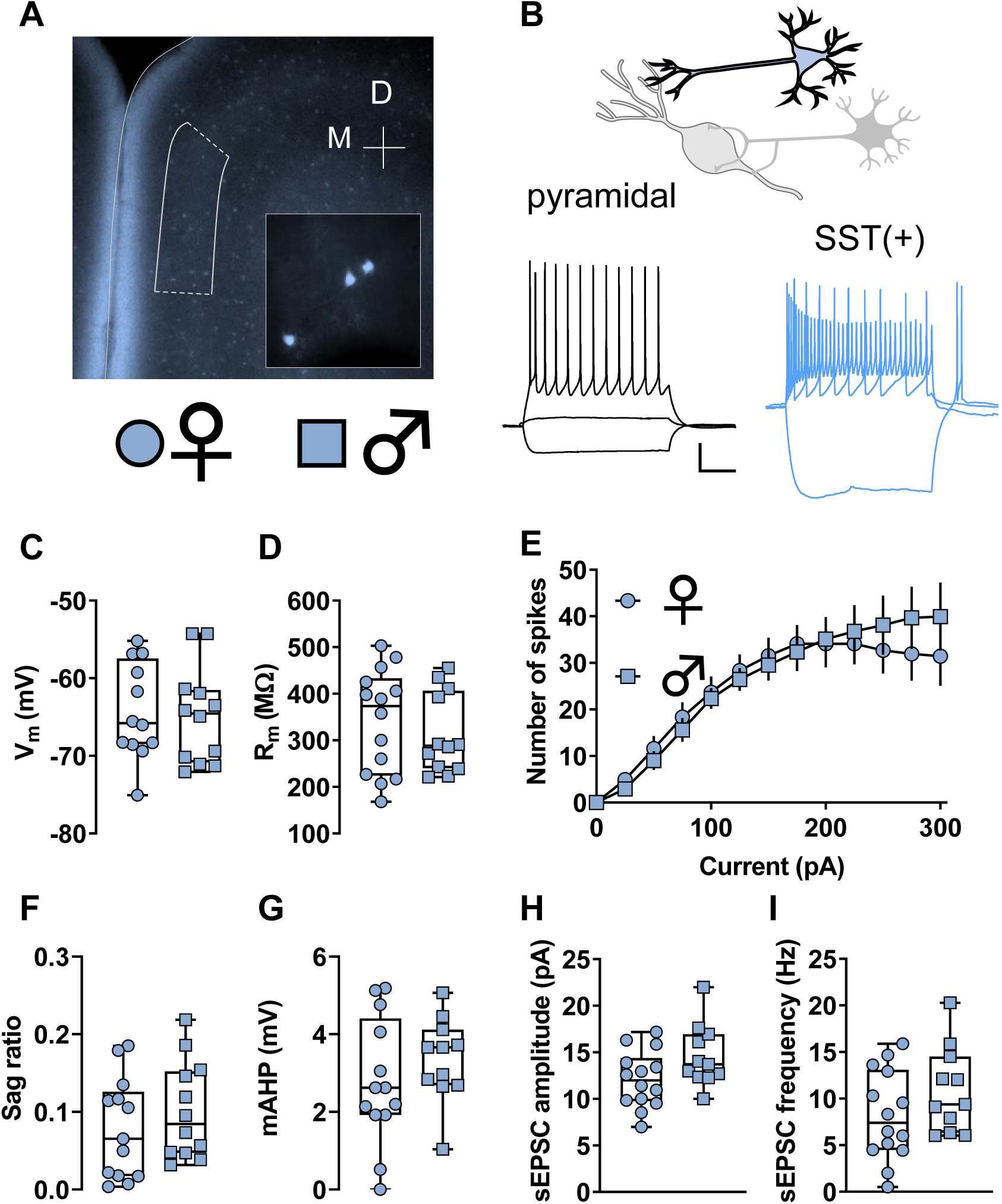
Somatostatin (SST)-(+) neurons display functional characteristics of Martinotti cells. **(A)** A representative 4X image displaying TdTomato fluorescence in deep players of the mouse prefrontal cortex (PFC). Boxed area indicates prelimbic subregion of the mouse PFC. Inset, 40X magnification image showing individual SST-expressing interneurons. M, medial; D, dorsal. **(B)** Whole-cell patch-clamp recordings were made from identified neurons in the PFC. Representative current-clamp recordings from an unlabeled pyramidal cell (left, black) and a TdTomato-labeled SST-IN (right, blue). The pyramidal cell displays modest input resistance (R_m_) and does not fire during 25 pA current injection current. In contrast, most SST-INs display high membrane resistance (R_m_) and fire action potentials in response to minimal current injection and hyperpolarization rebound. Scale bars indicate 20 mV and 250 ms. **(C)** Resting membrane potential (V_m_) in SST-INs from female mice (circles) and male mice (squares). n/N = 12/3 cells/mice per group. **(D)** No difference in R_m_ between SST-INs from female and male mice. n/N = 12/3, 14/3. **(E)** No difference in current-evoked spiking between SST-INs from female and male mice. n/N = 12/3. **(F)** No difference in hyperpolarization sag ratio between SST-INs from female and male mice. n/N = 13/3, 12/3. **(G)** No difference in medium afterhyperpolarization (mAHP) in SST-INs from female mice and male mice. n/N = 13/3. **(H)** No difference in spontaneous excitatory postsynaptic current (sEPSC) amplitude between SST-INs from female and male mice. n/N = 14/3, 11/3. **(I)** No difference in the mean of sEPSC frequency between SST-INs from female and male mice. n/N = 14/3, 11/3.

### 3.3 Adaptations to PV-IN and SST-IN membrane physiology in male, but not female, mice following IA ethanol

Across most rodent strains, females voluntarily consume more ethanol than males. This phenotype has been observed across many laboratories (Becker and Lopez, 2004; Hwa et al., 2011; Jury et al., 2017; McCall et al., 2013; Priddy et al., 2017), but the etiology underlying the sex difference remains incompletely understood. To model high levels of volitional alcohol drinking, we implemented an intermittent access (IA) schedule, where mice were provided with alternating days of ethanol availability in their home cages. Female mice drank more than males (Figure 1), and we then assessed intrinsic properties of PFC PV-INs and SST-INs at one-day abstinence from IA ethanol. We observed minimal effects of IA ethanol treatment on passive membrane properties or hyperpolarization sag in either interneuron subtype in either sex (Figure S1). Furthermore, we observed little effect of IA ethanol exposure on PV-IN active membrane properties in female mice (Figure 4A and 4B). In contrast, PV-INs from IA ethanol male mice exhibited distinct adaptations surrounding action potential initiation. Across multiple current injections, we observed increased current-evoked firing in PV-INs from male mice (Figure 4C). Following trains of action potentials, many neuron types display mAHP, a brief (∼100-ms to 2-sec) hyperpolarization. mAHP is generally mediated by voltage-and/or calcium-gated potassium channels and serves a feedback mechanism to limit continuous neural activity. PV-INs from male IA ethanol mice displayed greater mAHP than matched controls (Figure 4C). Together, these data indicate that PV-INs from male mice are hyperexcitable following IA ethanol and susceptible to enhanced feedback to limit their ongoing activity. We next examined membrane physiology of SST-INs. Like the PV-INs, SST-INs from female IA ethanol mice displayed similar current-evoked firing and mAHP to controls (Figure 4E and 4F). Unlike PV-INs, SST-IN spike-firing was no different between control and IA ethanol male mice (Figure 4G), but SST-INs from male IA ethanol mice did display enhanced mAHP (Figure 4H). Importantly, however, the large mAHP in male IA ethanol SST-INs might stem from a non-specific increase in Rm (Figure S1K) and therefore may not reflect a specific change to calcium-activated potassium channels. Overall, these intrinsic physiology adaptations suggest that IA ethanol alters the processes through which interneurons respond to synaptic input and alter PFC microcircuit function. We therefore aimed to better understand how binge drinking might modify excitatory input onto PFC PV-INs and SST-INs and investigated synaptic properties from controls and IA ethanol-exposed mice.

**Figure 4.**
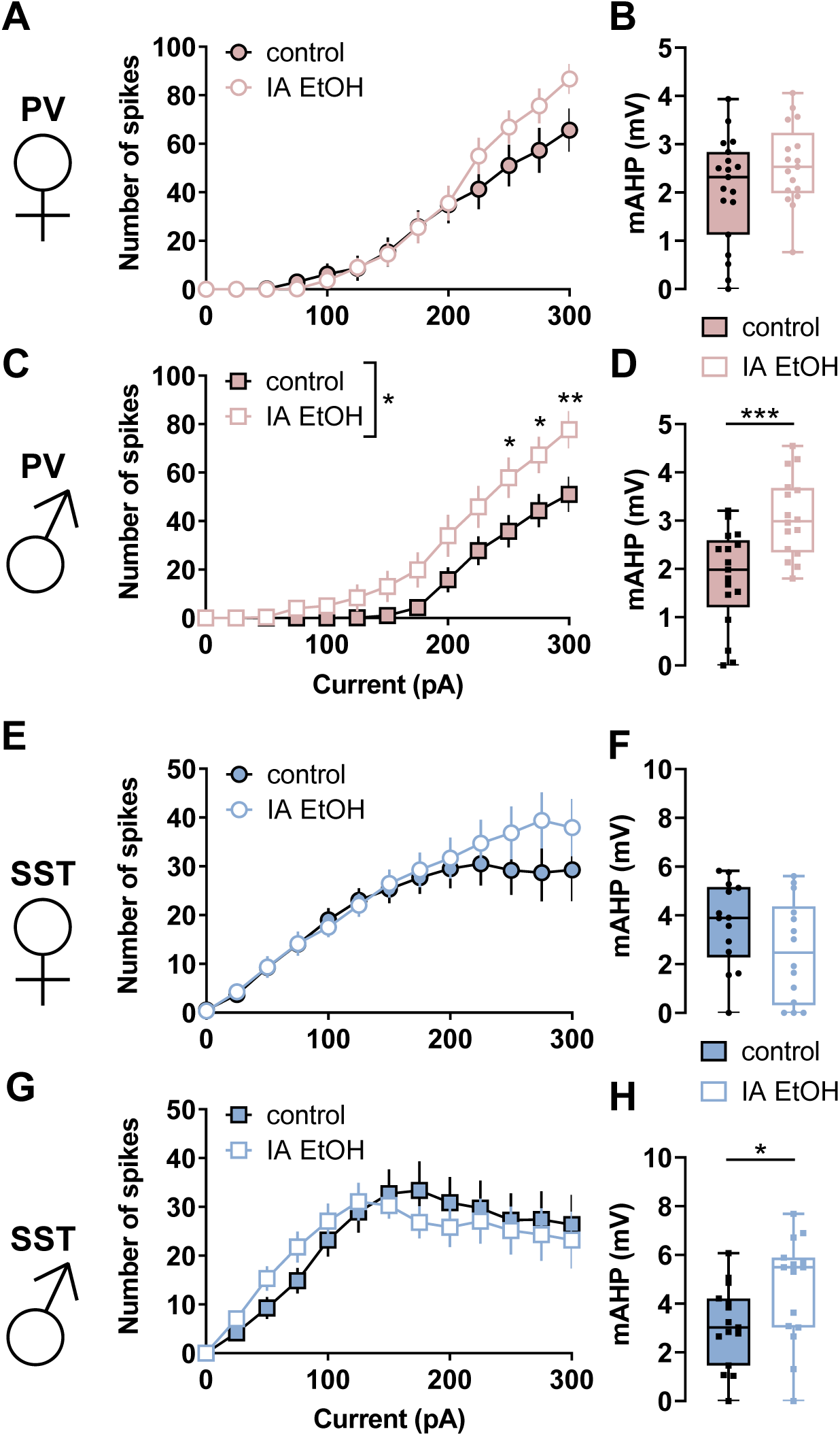
Intermittent ethanol increases medium afterhyperpolarization (mAHP) in parvalbumin (PV-IN) and somatostatin (SST-IN) interneurons in male mice. **(A)** Modest increase in spiking during large current injections in PV-INs from IA ethanol treated female mice relative to controls. (Two-way RM ANOVA main effect of input: F_12,432_ = 88.44, p < 0.0001; main effect of IA ethanol: F_1,36_ = 0.7002, n.s.; input x IA ethanol interaction: F_12,432_ = 2.213, p < 0.02). n/N = 17-19/5 cells/mice per group. **(B)** No difference in mAHP in IA ethanol PV-INs relative to controls in female mice n/N = 17-19/5. **(C)** Increased spiking in response to current injections in PV-INs from IA ethanol treated male mice relative to controls. (Two-way RM ANOVA main effect of input: F_12,348_ = 68.39, p < 0.0001; main effect of IA ethanol: F_1,29_ = 6.138, p < 0.02; input x IA ethanol interaction: F_12,348_ = 2.796, p < 0.002; *: p < 0.05, **: p < 0.01, Bonferonni post-tests). n/N = 16-17/4. **(D)** Enhanced mAHP in IA ethanol-treated PV-INs relative to controls in male mice (3.08 ± 0.21 vs 1.82 ± 0.24 mV, ***: p < 0.001, t-test). n/N = 16-17/5. **(E-F)** In female mice, no differences were observed in SST-IN current-evoked firing or mAHP between control (filled circles) and IA ethanol (open circles) treatment groups n/N = 13-20/4. **(G)** No difference in current-evoked spiking in SST-INs from IA ethanol treated male mice relative to controls. n/N = 16-17/4. **(H)** Increased mAHP in IA ethanol SST-INs relative to controls in male mice (3.08 ± 0.21 vs 1.82 ± 0.24 mV, ***: p < 0.001, t-test). n/N = 16-17/4.

### 3.4 Opposing changes to synaptic strength onto PFC PV-INs after IA ethanol in female and male mice

PFC interneurons generally display low levels of basal activity *in vivo*. Instead, long-range excitatory afferents dynamically recruit PV-IN and SST-IN activity to shape local networks of PFC pyramidal cells through feedforward and feedback inhibition. Thus, assessing how IA ethanol alters the synaptic strength of excitatory synapses onto PFC interneurons is essential to place the observed membrane physiology changes within a holistic context. We first analyzed the amplitude and frequency of sEPSCs to assess quantal size and content of glutamate synapses onto PV-INs (Figure 5A). In female mice, we observed a decrease in both sEPSC amplitude (Figure 5B) and frequency (Figure 5C) between IA ethanol treatment and controls. These data suggest PV-INs in female mice display reduced AMPA receptor function and fewer detectable synapses after IA ethanol exposure. We next used electrical stimulation to evaluate the paired-pulse ratio (PPR), which is modulated by changes in neurotransmitter release probability. We observed no difference in PPR (Figure 5D), suggesting changes in presynaptic glutamate release do not play a major role in the adaptations following IA ethanol. In PV-INs of female mice, the overall changes are most consistent with attenuated postsynaptic AMPA receptor function and fewer detectable synapses. Opposing results were obtained in PV-INs from male mice. IA ethanol increased both sEPSC amplitude (Figure 5E and 5F) and frequency (Figure 5E and 5G) without affecting PPR (Figure 5H), suggesting enhanced function of postsynaptic AMPA receptors and an increase in the number of detectable excitatory synapses after IA ethanol exposure. While we observed likely postsynaptic changes in both groups, these striking findings indicate that the excitatory synapses onto PFC PV-INs undergo diametrically opposed changes in female and male mice. These synaptic adaptations are expected to attenuate and facilitate PV-IN recruitment by excitatory drive in female and male mice with a history of IA ethanol respectively.

**Figure 5.**
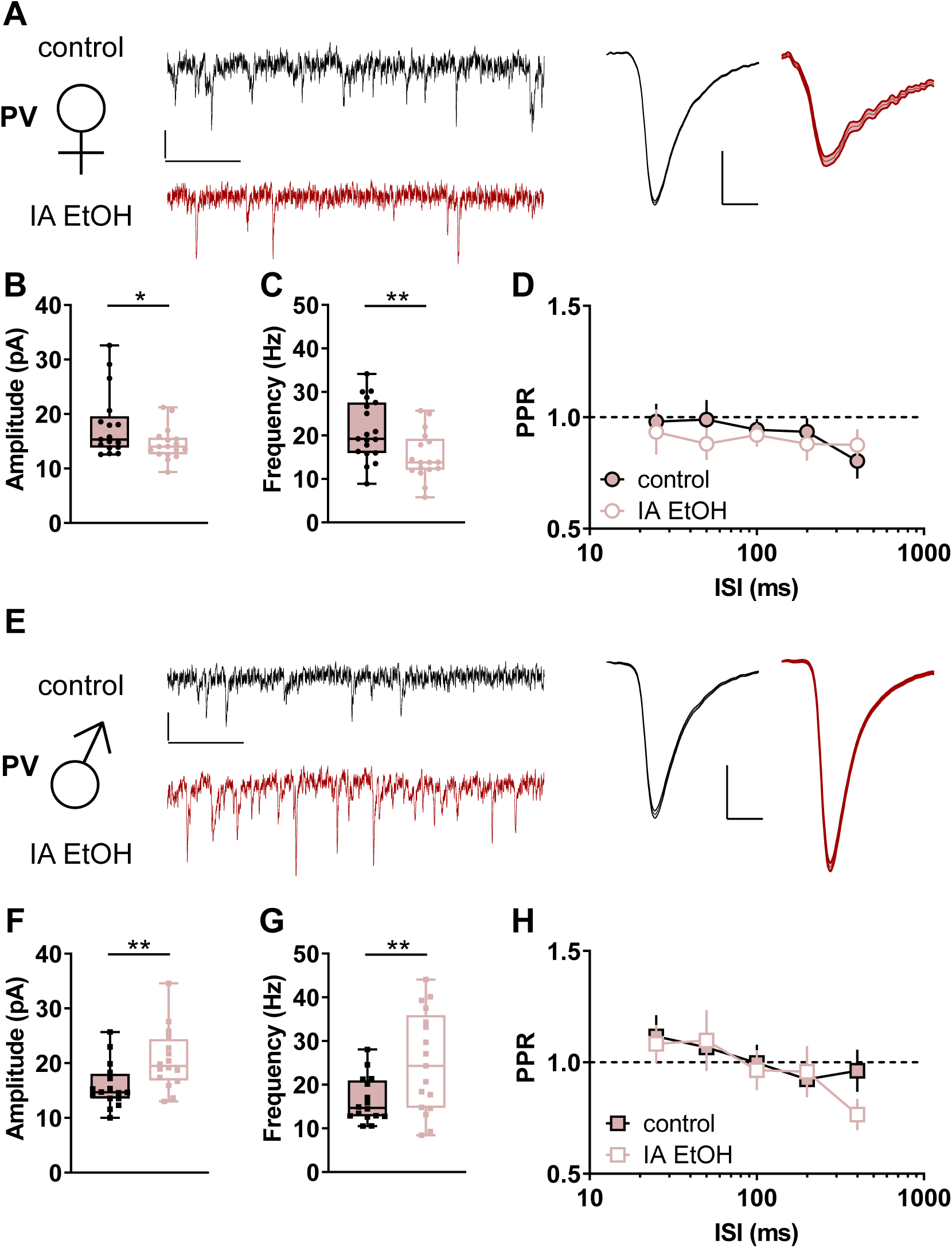
Intermittent access to ethanol generates diametrically opposing changes to excitatory synaptic strength onto parvalbumin-expressing interneurons (PV-INs) based on sex. **(A)** Left, Representative spontaneous excitatory postsynaptic current (sEPSC) traces from PV-INs from control (black) and IA ethanol (red) female mice. Scale bars indicate 10 pA, 100 ms. Right, Averaged sEPSC from representative experiment. Scale bars indicate 5 pA, 2 ms. **(B)** Decreased sEPSC amplitude in PV-INs from control and IA ethanol female mice. (14.5 ± 0.7 vs 17.9 ± 1.5 pA, *: p < 0.05, t-test) n/N = 17/5 cells/mice per group **(C)** PV-INs from female mice given IA ethanol exhibited decreased sEPSC frequency relative to controls (15.2 ± 1.3 vs 21.2 ± 1.6 Hz, **: p < 0.01, t-test). n/N = 17-19/5. **(D)** No group difference in paired-pulse ratio (PPR) between control and IA ethanol female mice across several interstimulus intervals (ISIs). n/N = 7-9/3. **(E)** Left, Representative sEPSC traces from PV-INs from control (black) and IA ethanol (red) male mice. Scale bars indicate 10 pA, 100 ms. Right, Averaged sEPSC from representative experiment. Scale bars indicate 5 pA, 2 ms. **(F)** Greater sEPSC amplitude in PV-INs from IA ethanol group relative to control male mice (20.7 ± 1.4 vs 15.9 ± 1.0 pA, **: p < 0.01, t-test). n/N = 16/5. **(G)** PV-INs from male mice given IA ethanol displayed increased sEPSC frequency relative to controls (25.1 ± 2.8 vs 16.5 ± 1.3 Hz, **: p < 0.01, t-test). n/N = 16-17/5. **(H)** No group difference in PPR between control and IA ethanol male mice. n/N = 7-9/3.

### 3.5 Diminished synaptic strength onto PFC SST-INs after IA ethanol

As with PV-INs, we evaluated changes to excitatory synaptic transmission onto SST-INs in control mice and those exposed to IA ethanol. In female mice, one-day abstinence from IA ethanol was associated with decreased excitatory synaptic strength (Figure 6A), as evidenced by reductions in both sEPSC amplitude (Figure 6B) and frequency (Figure 6C). The PPR of evoked EPSCs was not different between the control and IA ethanol groups, suggesting these changes in excitatory transmission occurred through postsynaptic reduction in AMPA receptor function and the number of detectable synapses. We observed similar changes in male mice (Figure 6E). IA ethanol produced an attenuation of sEPSC amplitude (Figure 6F) and frequency (Figure 6G) in SST-INs of male mice. Surprisingly, we observed decreased PPR across multiple interstimulus intervals in these cells from the IA ethanol group (Figure 6H). At face value, these data suggest increased presynaptic glutamate release probability and are not consistent with the concomitant reduction in sEPSC frequency. We offer two potential explanations for this discrepancy: (1) distinct sets of synapses may have been sampled during spontaneous and evoked EPSC recordings, as has been observed at excitatory synapses onto other cell types in the central nervous system (Ramirez and Kavalali, 2011); and (2) SST-IN PPR may be regulated by a postsynaptic feature, such as the activity-dependent polyamine sensitivity of AMPA receptors previously observed in cortical interneurons (Rozov and Burnashev, 1999). Nonetheless, the collective dataset suggests that IA ethanol exposure attenuates excitatory transmission overall onto SST-INs in all groups of mice.

**Figure 6.**
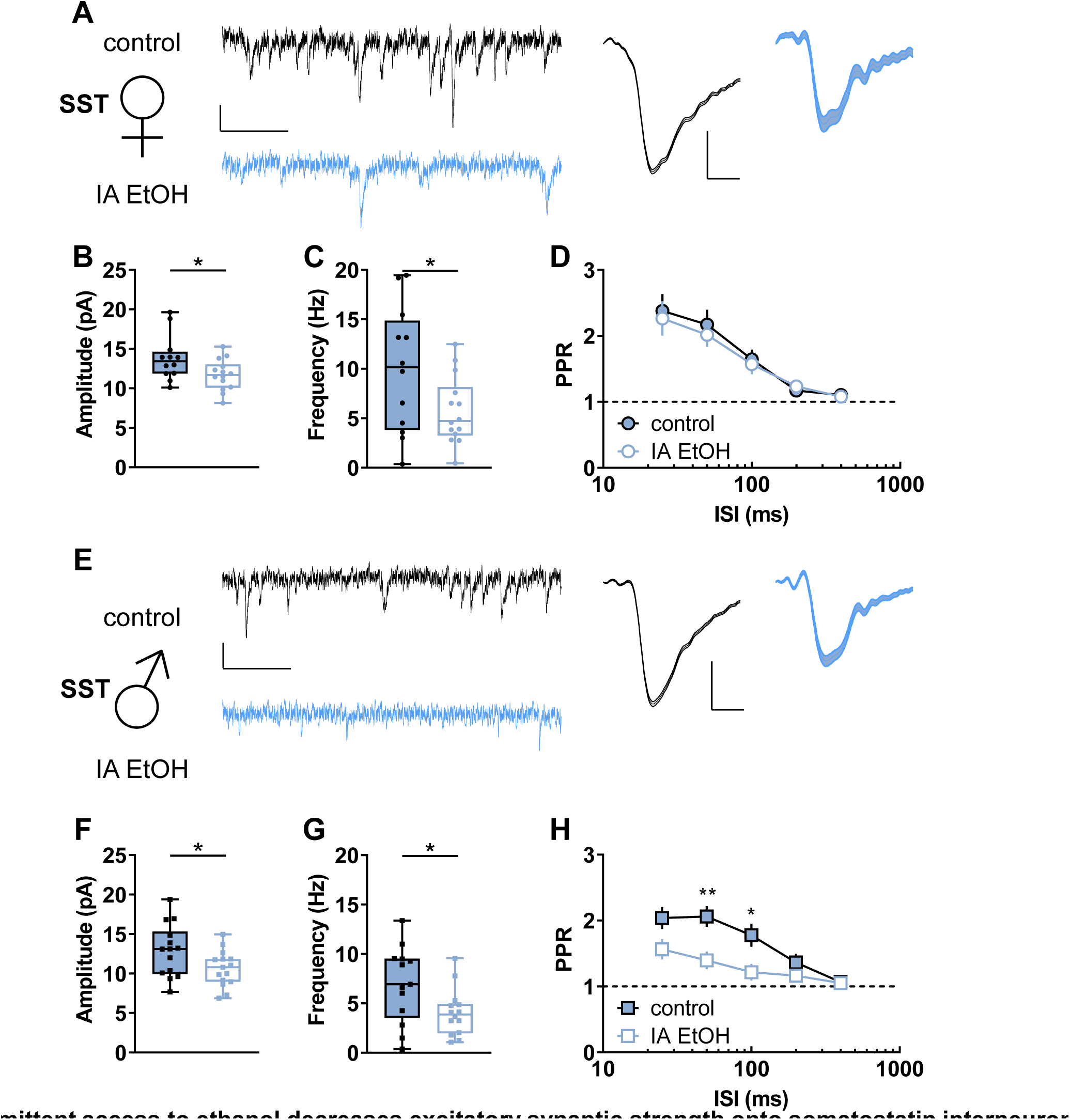
Intermittent access to ethanol decreases excitatory synaptic strength onto somatostatin interneurons (SST-INs). **(A)** Left, Representative spontaneous excitatory postsynaptic current (sEPSC) traces from SST-INs from control (black) and IA ethanol (blue) female mice. Scale bars indicate 10 pA, 100 ms. Right, Averaged sEPSC from representative experiment. Scale bars indicate 5 pA, 2 ms. **(B)** Decreased sEPSC amplitude was observed in SST-INs from IA ethanol female mice relative to controls. controls (11.7 ± 0.5 vs 13.8 ± 0.8 pA, *: p < 0.05, t-test). n/N = 12-14/4 cells/mice per group. **(C)** SST-INs from female mice given IA ethanol exhibited decreased sEPSC frequency relative to controls (5.7 ± 0.9 vs 9.9 ± 1.8 Hz, *: p < 0.05, t-test). n/N = 12-19/4-5 cells/mice. **(D)** SST-IN paired-pulse ratio (PPR) did not differ between control and IA ethanol female mice across multiple interstimulus intervals (ISIs). n/N = 9-11/3-4. **(E)** Left, Representative sEPSC traces from SST-INs from control (black) and IA ethanol (blue) male mice. Scale bars indicate 10 pA, 100 ms. Right, Averaged sEPSC from representative experiment. Scale bars indicate 5 pA, 2 ms. **(F)** Decreased sEPSC amplitude in SST-INs from IA ethanol group relative to control male mice (10.6 ± 0.6 vs 12.9 ± 0.9 pA, *: p < 0.05, t-test). n/N = 14-15/5. **(G)** SST-INs from male mice given IA ethanol displayed decreased sEPSC frequency relative to controls (4.1 ± 0.6 vs 6.9 ± 1.1 Hz, *: p < 0.05, t-test). n/N = 16-17/5. **(H)** Decreased PPR in IA ethanol male mice (Two-way RM ANOVA main effect of ISI: F_4,84_ = 24.85, p < 0.0001; main effect of IA ethanol: F_1,21_ = 6.897, p < 0.02; input x IA ethanol interaction: F_12,84_ = 4.6, p < 0.003; *: p < 0.05, **: p < 0.01, Bonferonni post-tests) n/N = 11-12/4-5.

## 4. Discussion

The PFC is essential for top-down moderation of drinking (Abernathy et al., 2010; George and Koob, 2010). The preclinical literature has primarily focused on how ethanol alters the function of PFC pyramidal cells, the principal neurons that convey information from the PFC to subcortical structures. Nonetheless, PFC output is dynamically regulated by local interneurons, so we sought to investigate physiological adaptations occurring on those discrete cell types following ethanol exposure. We functionally interrogated two genetically defined subtypes of PFC interneurons in a binge drinking model. IA ethanol induced disparate sex-dependent adaptations to PV-INs – fewer detectable synapses in female mice and enhanced postsynaptic AMPA receptor function in male mice. In contrast, IA ethanol dampened excitatory synaptic strength onto SST-INs in all groups of mice. Together, these data indicate that altered excitatory synaptic drive onto PV-INs and SST-INs may contribute to PFC dysfunction in AUD.

The changes we observed to PV-IN and SST-IN intrinsic and synaptic physiology may or may not be related. While we observed a significant increase in mAHP following IA ethanol in interneurons from male mice, a potential increase may have been occluded in PV-INs from female mice due to their relatively large basal mAHP. Following sustained depolarization, the mAHP limits neural activity and calcium mobilization within a postsynaptic cell. Therefore, the coincidental ethanol-induced increases in PV-IN mAHP and synaptic strength in male mice are consistent with a homeostatic response, i.e. increased mAHP might provide a feedback mechanism to mitigate excessive activation of PV-INs following synaptic stimulation. In PV-INs from female mice and SST-INs, by contrast, an IA ethanol-induced increase in mAHP might have facilitated spike timing-dependent long-term depression (Lu et al., 2007), thereby initiating an ethanol-induced reduction synaptic strength. In addition – or as an alternative – to these potential causal relationships, the coincidental changes to mAHP and synaptic strength might be manifested by the further separation of PV-INs into responsive and non-responsive subpopulations. PV-INs can be divided into “basket cells” and “chandelier cells” based on the subcellular targeting of their axons. Chandelier cells exhibit minimal mAHP relative to basket cells (Povysheva et al., 2013), thus one intriguing hypothesis is that IA ethanol specifically enhances mAHP (and also modulates synaptic strength) in this PV-IN subtype. Similarly, cortical SST-INs can be further stratified into subclasses based on intrinsic properties, connectivity, and protein expression (Tremblay et al., 2016), and some of these factors may confer susceptibility to alcohol-related pathophysiology.

Female mice drink more ethanol than male counterparts. While that could conceivably contribute to differences in our observed physiological changes, we find it unlikely that relatively modest variation in ethanol exposure initiates fundamentally distinct changes in physiology.

With that in mind, the current findings reveal that PFC PV-INs undergo entirely distinct adaptations based on sex, particularly with regards to synaptic physiology. These results beg the question, do ethanol-induced adaptations to PV-INs contribute to sex differences in drinking behaviors? Substantial further research is needed to address this question. For one, it remains unclear how PFC PV-IN activity *in vivo* modulates drinking and other appetitive behaviors.

Experiments designed to monitor and manipulate PV-IN activity during a variety of components of ethanol-seeking tasks will be important to fully understand how molecular changes to PV-IN physiology confer AUD-like adaptations. One overly simple and wildly speculative prediction is that feedforward drive onto PV-INs inhibits cortical circuits involved in drinking behaviors, such as the PFC projections to the nucleus accumbens (Seif et al., 2013) or periaqueductal gray (Siciliano et al., 2019). Were that the case, decreased excitatory transmission onto PV-INs could facilitate drinking in female mice and enhanced drive onto PV-INs in male mice might confer some resilience. Similarly, attenuated drive onto SST-INs might promote drinking in all groups of mice, but the discrepant changes to PPR and sEPSC frequency in male SST-INs highlight the need to investigate specific sources of glutamate onto PFC interneurons. In addition to providing top-down control over drinking, the PFC regulates a variety of cognitive processes with relevance for AUD. In particular, working memory and cognitive flexibility are regulated by PFC PV-INs (Murray et al., 2015) and SST-INs (Abbas et al., 2018). Based on this, we predict that interneuron dysfunction likely contributes to impairments in PFC-dependent cognitive processes observed in animal models of AUD (George et al., 2012; Salling et al., 2018; Trantham-Davidson et al., 2014; Vargas et al., 2014). Further research, however, is needed to test this hypothesis and to examine potential sex differences in AUD-related cognitive disruptions.

The preclinical literature modeling PFC dysfunction has generally utilized male subjects and focused on adaptations to pyramidal cells. Studies describing PFC interneuron dysfunction have been more limited and exclusively examined the physiological ramifications of ethanol dependence or chronic intoxication. Trantham-Davidson et al. (Trantham-Davidson et al., 2014) assessed intrinsic properties of functionally identified fast-spiking interneurons (putative PV-INs) in male rats exposed to CIE. CIE generated differences in dopamine modulation of fast-spiking interneuron current-evoked firing, but baseline excitability parameters, including input-output firing curves and mAHP, were not reported. Another recent paper by Hughes et al. (Hughes et al., 2019) described several changes to the intrinsic properties of PFC interneurons in a rat model of chronic intoxication. Following ethanol exposure, Hughes et al. observed decreased fast-spiking interneuron excitability in all rats, while we detected *increased* PV-IN spiking in male mice only. Further, Hughes et al. observed ethanol-induced sex differences in current-evoked firing in putative Martinotti cells, while we noted minimal effects on SST-IN intrinsic physiology in either female mice or male mice. Several major technical differences are likely to explain these discrepancies. Hughes et al. delivered an intoxicating dose of ethanol (∼250 mg/dL) to rats on successive days, whereas mice in the current studies voluntarily drank on alternating nights to reach moderate blood ethanol concentrations (∼80 mg/dL reported in previous studies (Hwa et al., 2011; Salling et al., 2018)). In addition to methodological differences in slice preparation and interneuron classification, differences between the two studies might stem from means of ethanol delivery, overall intake, pattern of intake, and species. Each of these parameters merits further examination. We believe the diversity of preclinical AUD models is a beneficial feature of the research community; continued research across several models well help elucidate the core features of disease etiology. To our knowledge, the present study represents the first characterization of PFC interneuron synaptic physiology following voluntary drinking, although these findings are also limited by a single timepoint. Future studies should be designed to address the development and recovery of PFC interneuron pathophysiology and related behavioral adaptations across multiple disease models.

Based on the current findings, one would expect for inhibitory transmission onto PFC pyramidal cells to be altered following chronic drinking and in models of alcohol dependence. In ostensible contrast to that hypothesis, previous research did not reveal differences in inhibitory transmission onto deep layer prelimbic PFC pyramidal cells during acute withdrawal from CIE (Pleil et al., 2015; Trantham-Davidson et al., 2014). During those experiments, however, GABAergic inhibitory postsynaptic currents (IPSCs) on pyramidal cells were collected with non-specific electrical stimulation or in a spontaneous manner. The present findings suggest that output from each interneuron subpopulation is likely altered following ethanol exposure, but coincidental adaptations might have obfuscated changes in non-specific IPSCs in previous work. Our findings suggest this possibility is particularly likely for male subjects, as we observed contrasting changes to synaptic strength onto PV-INs and SST-INs in male mice. Consistent with this hypothesis, Trantham-Davidson et al (Trantham-Davidson et al., 2014). discovered that CIE disrupted the ability of D4 dopamine receptors to modulate IPSCs recorded from pyramidal cells. D4 receptors regulate PV-IN function in the PFC (Zhong and Yan, 2016) and hippocampus (Andersson et al., 2012) without affecting other types of interneurons, suggesting that PV-INs mediated the CIE-induced changes to D4 signaling observed by Trantham-Davidson et al (2014). Future investigations targeting PFC inhibitory synapses with interneuron type-specific manipulations, in both female and male rodents, should improve our understanding of how alcohol exposure dysregulates PFC function and assist efforts to discover novel targets for the treatment of AUD.

Restoring normal synaptic physiology on PFC PV-INs and SST-INs following IA ethanol may provide avenues to mitigating behavioral disruptions relevant to AUD. A better understanding of the mechanisms regulating synaptic plasticity on cortical interneurons would be essential towards these endeavors. Calcium-permeable AMPA receptors and NMDA receptors represent two candidate molecules that might modulate synaptic strength onto PV-INs and SST-INs during or after IA ethanol. Future studies should therefore examine drinking-related changes in AMPA and NMDA receptor expression, stoichiometry, and signaling on cortical interneurons. In addition to ionotropic glutamate receptors, changes in fast glutamate transmission are often mediated by metabotropic glutamate (mGlu) receptors. Several previous findings suggest mGlu receptors may be involved in the PFC interneuron synaptic pathophysiology we observed after IA ethanol. Transcript and protein for mGlu receptor subtypes 1 (mGlu1) and 5 (mGlu5) are enriched in interneurons, and mGlu1 and mGlu5 have been implicated in forms of long-term potentiation specific to interneurons (Le Duigou and Kullmann, 2011; Perez et al., 2001). Furthermore, mGlu5, but not mGlu1, gates long-term potentiation onto fast-spiking interneurons in the visual cortex (Sarihi et al., 2008), but these plasticity mechanisms have not, to our knowledge, been investigated in the PFC or other associative cortices. In addition to these plasticity mechanisms, pronounced sex differences in mGlu1 and mGlu5 signaling have been observed in the limbic system: estradiol regulates hippocampal inhibitory transmission through mGlu1 (Huang and Woolley, 2012), and sex hormones modulate mGlu5-dependent synaptic plasticity in the nucleus accumbens (Gross et al., 2018; Peterson et al., 2015). Altogether, the breadth of preclinical literature raises the possibility that changes in PFC glutamate transmission during IA ethanol might recruit mGlu1/mGlu5 signaling to modulate synaptic strength on PV-INs and SST-INs in a sex-dependent manner. Consistent with that hypothesis, mice harboring mGlu5 ablation from PV-expressing neurons display altered sensitivity to rewarding drugs and a sex-dependent increase in habitual behavior (Barnes et al., 2015). Finally, small molecule modulators of mGlu1 and mGlu5 are efficacious in multiple preclinical AUD models (Joffe et al., 2018), providing further impetus for investigating how these signaling pathways may underlie disease etiology.

## 5. Conclusion

Interneurons arise from distinct progenitors and express distinct transcriptional program, presenting opportunities to leverage biological idiosyncrasies for the targeted treatment of disease. Efficacious AUD treatments might one day be developed from cortical interneuron dysfunction initially observed in a preclinical disease model. The striking adaptations to synaptic physiology described here provide one potential starting place.

## Acknowledgements

The authors thank members of the Conn and Winder labs for stimulating discussions. This work was supported by National Institutes of Health grants R01MH062646 and R37NS031373 (P.J.C.). M.E.J. was supported by a postdoctoral fellowship through the Pharmaceutical Research and Manufacturers of America Foundation.

## Author Contributions

Conceptualization, M.E.J; Investigation, M.E.J.; Supervision, D.G.W. and P.J.C.; Writing – Original Draft, M.E.J.; Writing – Review & Editing – all authors; Funding Acquisition, M.E.J. and P.J.C.

---

**Figure S1.**
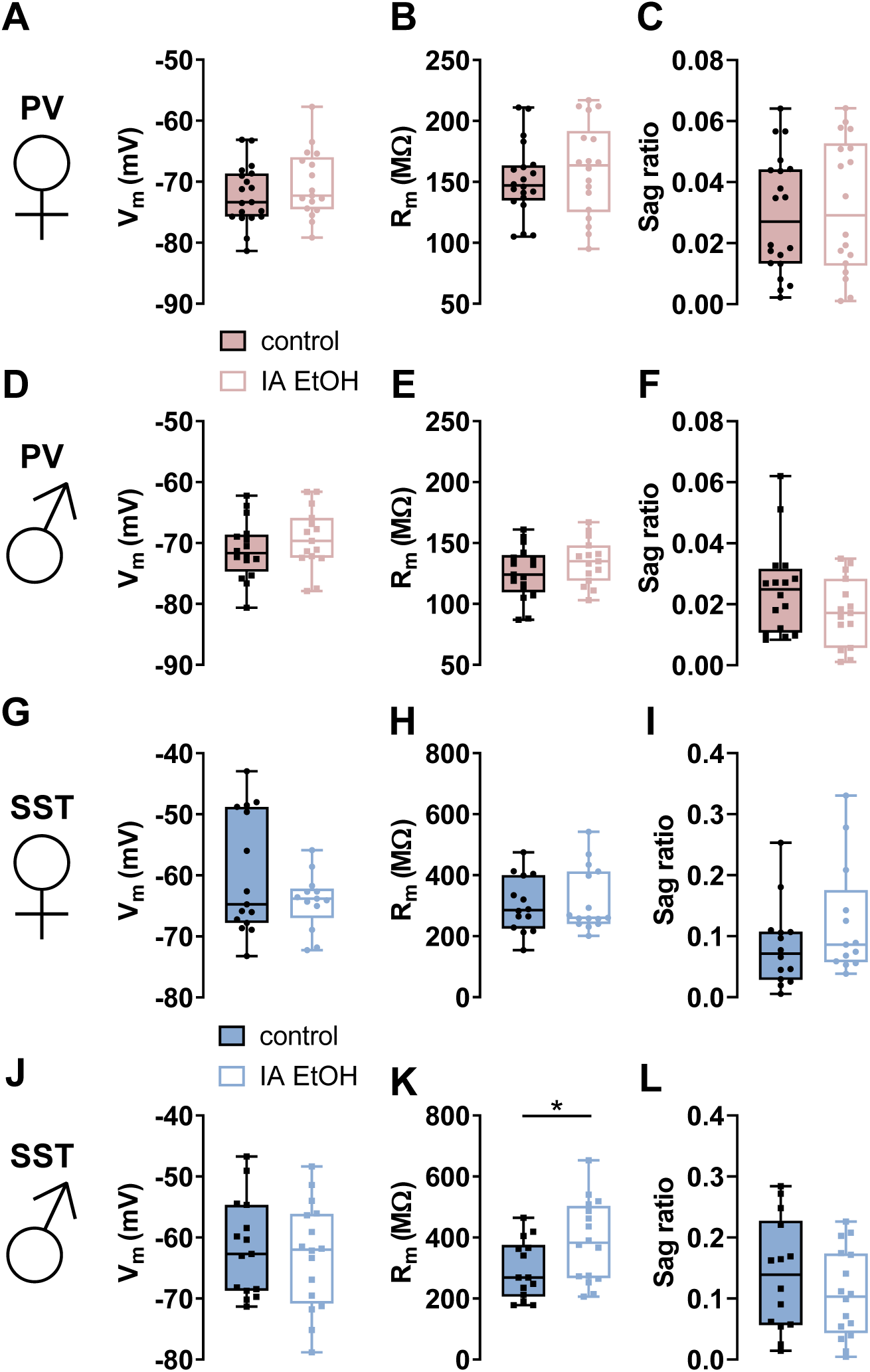
Minimal adaptations to several interneuron membrane properties following intermittent access to ethanol. **(A-C)** In female mice, no differences were observed in parvalbumin interneuron (PV-IN) resting membrane potential (V_m_), membrane resistance (R_m_), or sag ratio between control (filled circles) and IA ethanol (open circles) treatments. n/N = 17-20/5 cells/mice per group. **(D-F)** In male mice, no differences in PV-IN V_m_, R_m_, or sag ratio were detected between control (filled squares) and IA ethanol (open squares) treatments. n/N = 15-17/4. **(G-I)** In female mice, no differences were observed in somatostatin interneuron (SST-IN) V_m_, R_m_, or sag ratio between control and IA ethanol treatments. n/N = 13-20/4-5. **(J)** In male mice, SST-IN V_m_ did not differ between control and IA ethanol treatments. n/N = 15-17/4. **(K)** IA ethanol was associated with increased R_m_ in SST-INs from male mice (389.1 ± 33.3 vs 295.6 ± 25.7 MΩ, *: p < 0.05, t-test). n/N = 14-16/4. **(L)** No difference in SST-IN sag ratio between control and IA ethanol male mice. n/N = 14-16/4.

